# Extending island biogeography theory to biotic islands: Microbial communities in epiphytic bird’s nest fern *Asplenium nidus*

**DOI:** 10.64898/2026.03.30.715435

**Authors:** Yu-Pei Tseng, Shuo Wei, Po-Ju Ke

**Affiliations:** Institute of Ecology and Evolutionary Biology, National Taiwan University, Taipei, Taiwan; School of the Environment, The University of Queensland, Brisbane, Australia; Yale School of the Environment, Yale University, New Haven, Connecticut, USA

**Author notes:** Correspondence author: Institute of Ecology and Evolutionary Biology, National Taiwan University, Taipei 10617, Taiwan.

**Keywords:** Species–area relationship, metacommunity theory, environmental heterogeneity, biotic insular system, distance-decay

## Abstract

1. Biotic insular systems differ from conventional islands because patch attributes change dynamically as patch-forming organisms develop. It therefore remains unclear whether the assembly mechanisms predicted by island biogeography theory (IBT) operate in such systems. Here, using epiphytic bird’s nest ferns (BNFs, *Asplenium nidus*) as a model biotic island system, we tested whether fungal and bacterial community diversity conform to species–area relationships predicted by IBT. With a stratified sampling scheme, we further evaluated the underlying mechanisms (passive sampling, disproportionate effects, and environmental heterogeneity) of species–area relationships, and assessed isolation effects using distance–decay patterns in community similarity.

2. We treated each BNF individual as a microbial island and categorized 24 BNFs into three size classes. Microbial and humus samples from multiple litter layers within each BNF individual were collected; microbial communities were characterized using next-generation sequencing, and humus chemical properties (pH and C:N ratio) were measured to characterize microhabitat conditions. To investigate mechanisms underlying species–area relationships, we applied a multi-scale rarefaction framework to partition diversity components. Spatial distances among BNFs were quantified to evaluate isolation effects.

3. Consistent with IBT predictions, both fungal and bacterial communities exhibited positive species–area relationships, indicating that larger BNFs harbored greater microbial richness. Diversity partitioning suggested that fungal richness increased through both disproportionate effects and environmental heterogeneity, whereas bacterial richness was primarily driven by environmental heterogeneity. Within larger ferns, greater heterogeneity in litter pH was associated with increased species turnover across litter layers, suggesting that decomposition-driven pH gradients create diverse microhabitats that promote microbial diversity. In addition, both microbial communities exhibited distance–decay patterns, indicating that isolation contributes to community assembly through dispersal limitation.

**4. Synthesis.** Our results demonstrate that BNFs function as a biotic insular system, in which both patch size and spatial isolation structure microbial diversity, consistent with predictions from IBT. Furthermore, we show that environmental heterogeneity generated by the growth of the habitatforming BNF mechanistically links island area to microbial diversity. Our study integrates both local habitat heterogeneity and regional spatial structure, highlighting the potential to extend IBT and metacommunity theory to organism-formed habitats.

## Introduction

Ecological communities are shaped by both local processes, such as species interactions and environmental heterogeneity, and regional processes that regulate colonization and extinction through dispersal (MacArthur & Levins, 1967, Chesson, 1986, Wootton, 1994). Regional processes become a key determinant of community assembly when local communities are connected by dispersal, motivating the development of spatial ecological theories that explicitly consider area, isolation, and connectivity (Shurin & Allen, 2001, Hillebrand & Blenckner, 2002). Among these, the island biogeography theory provides foundational predictions linking island area and isolation to species richness via colonization–extinction dynamics. Insular habitats, which are discrete, island-like patches embedded within a surrounding matrix, have therefore served as model systems for testing these ideas. While empirical tests of island biogeography originally focused on abiotic insular systems, such as oceanic islands or habitat fragments, biotic insular systems may also function as discrete “biotic islands” that host diverse communities (Drakare *et al*., 2006, Matthews *et al*., 2016).

Biotic insular systems are found in many ecosystems and are defined as habitat patches formed, modified, and maintained by organisms themselves (Miller *et al*., 2018, Itescu, 2019). In these systems, island area, habitat heterogeneity, and extinction risk may change dynamically due to the development and biotic feedbacks of the patch-forming organism, potentially altering the mechanisms underlying classical island biogeography predictions. Although biotic islands have received increasing attention, empirical studies report mixed support of classical island biogeography patterns, with some systems conforming to area and isolation effects (e.g., Peay *et al*., 2007, Adams *et al*., 2017) while others deviating from them (e.g., Andrews *et al*., 1987, Kinkel *et al*., 1987). Consequently, it remains unclear how classical island biogeography predictions apply to biotic insular systems, and how regional processes associated with area and isolation interact with organism-mediated habitat dynamics within these systems.

One of the central predictions of island biogeography is the island species–area relationship (ISAR), which posits that species richness increases with the island’s size through three main mechanisms: passive sampling, disproportionate effects, and habitat heterogeneity. *Passive sampling* refers to the null expectation that larger patches sample a larger proportion of individuals from the regional species pool, thereby increasing the likelihood of finding more species in these larger patches (Connor & McCoy, 1979). The *disproportionate effect* highlights that certain ecological processes may affect species differently depending on patch size. For example, smaller patches often support smaller populations, which are more vulnerable to stochastic fluctuations, potentially leading to higher extinction risks and lower species richness (Chase *et al*., 2019). Finally, *habitat heterogeneity* posits that larger patches contain more varied microhabitats, offering diverse ecological niches that support a greater variety of species (Williams, 1964, Currie, 1991, Rosenzweig, 1995, MacArthur & Wilson, 1967). Despite solid empirical support for these mechanisms in abiotic insular systems, far fewer studies have tested whether they operate in biotic insular systems, where the three mechanisms may be tightly coupled through organism-driven processes. Moreover, because studies in biotic insular systems have often focused on local-scale assembly processes (e.g., Dinnage *et al*., 2019), it remains unclear whether these systems conform to the regional-scale predictions of island biogeography theory.

In addition to area, another key spatial factor in island biogeography theory is isolation, which historically refers to an island’s distance from the mainland species source. Islands further from the species source typically exhibit lower species richness due to reduced immigration, while islands closer to the species source tend to experience higher immigration and therefore higher species richness (MacArthur & Wilson, 1967). This concept extends naturally to metacommunity theory (Gilpin & Hanski, 1991, Wilson, 1992, Leibold *et al*., 2004), which extends island biogeography theory by emphasizing how dispersal among patches interacts with local ecological processes to shape community composition. In this context, neighboring communities rather than a mainland pool act as the propagule source (Fukami, 2005), and geographic distance among patches serves as a proxy for isolation. Increasing distance may generate a distance-decay pattern in species composition similarity, such that communities become less similar as the distance between patches increases (Soininen *et al*., 2007). In this framework, dispersal limitation represents a primary regional mechanism underlying distance-decay patterns, as reduced movement of organisms among distant patches constrains colonization and promotes spatial differentiation in community composition.

Epiphytic bird’s nest ferns (*Asplenium nidus*) provide a natural biotic island system in forest canopies, making them an ideal model for testing whether classical island biogeography theory predictions extend to organism-formed habitats. Commonly found on tree trunks and larger branches in tropical and subtropical forests in the Old World (e.g., Hsu *et al*., 2002, Ellwood *et al*., 2002), they form discrete, yet potentially dispersal-connected, habitat patches that vary substantially in size and spatial isolation. Their rosette-forming fronds efficiently intercept litterfall from the canopy, and the accumulated litter, senesced fronds, and adventitious roots together form a nest-like suspended soil habitat. These arboreal nests function as distinct patches to host diverse communities of arthropods (Ellwood & Foster, 2004, Karasawa & Hĳii, 2006) and microbes (Donald *et al*., 2020).

Building on this biotic insular framework, we used bird’s nest ferns as a model system to test whether core predictions of island biogeography theory extend to biotic islands by examining how fern size and isolation shape fungal and bacterial communities within individual ferns. We treated each fern individual as a discrete island, with litter layers within ferns representing local habitat units. We hypothesized that increases in fern size, used as a proxy for island area, would be associated with higher microbial richness, consistent with classical ISAR. In addition to testing whether a species–area relationship emerges in this system, a central novelty of our study is that we explicitly tested how the three mechanisms underlying ISAR operate in this biotic insular system. Specifically, we evaluated the relative support for the three mechanisms — *passive sampling*, *disproportionate effects*, and *environmental heterogeneity* — using the multi-scale rarefaction framework of Chase *et al*. (2019), in which *γ*, *α*, and *β* diversity correspond to a single BNF individual, a layer within a BNF individual, and difference among layers, respectively. Given the organism-formed nature of this system, we further hypothesized that environmental heterogeneity would be a particularly important contributor, such that larger and older ferns, which retain more organic material across various stages of decomposition, would exhibit greater structural complexity. Accordingly, we aimed to identify which environmental factors associated with litter accumulation and decomposition most strongly contribute to within-patch heterogeneity and microbial community structure. Finally, we hypothesized that increasing isolation, quantified as inter-patch distance, would reduce immigration rates and generate distance–decay patterns in microbial community similarity, consistent with dispersal limitation. Together, these hypotheses allow us to evaluate whether core predictions of island biogeography extend to biotic island systems.

## Materials and Methods

### Study species and study site

We studied the biotic insular system formed by *Asplenium nidus* (bird’s nest ferns; hereafter BNF), a common epiphytic fern widely distributed in humid forests of Taiwan. Individuals of *A. nidus* occur in a wide range of sizes, reflecting their long lifespan (Tsai *et al*., 2023) — large individuals have fronds that extend to approximately 150 cm in length and form rosettes with a diameter up to 200 cm (Martin *et al*., 2004). Our study site is located at Tonghou Historical Trail near the Tonghou riverside, New Taipei City, Taiwan (24°83’65” N, 121°65’37” E; elevation 450-520 m a.s.l.). To minimize the effect of large-scale environmental heterogeneity (e.g., variation in litter input associated with different host tree species), all *A. nidus* individuals were collected in a *Cryptomeria japonica* plantation in April 2023 (Fig. 1A & B). The site receives high annual rainfall of about 3600 mm per year and retains high air humidity throughout the year (mean annual air humidity is 91.9%) without an obvious dry season (based on mean annual precipitation from 1990 to 2023, CODiS, https://codis.cwa.gov.tw/StationData). The mean annual temperature is 19.7 ℃, with the coldest month in January (13.1 ℃) and the hottest month in July (25.8 ℃).

**Figure 1.**
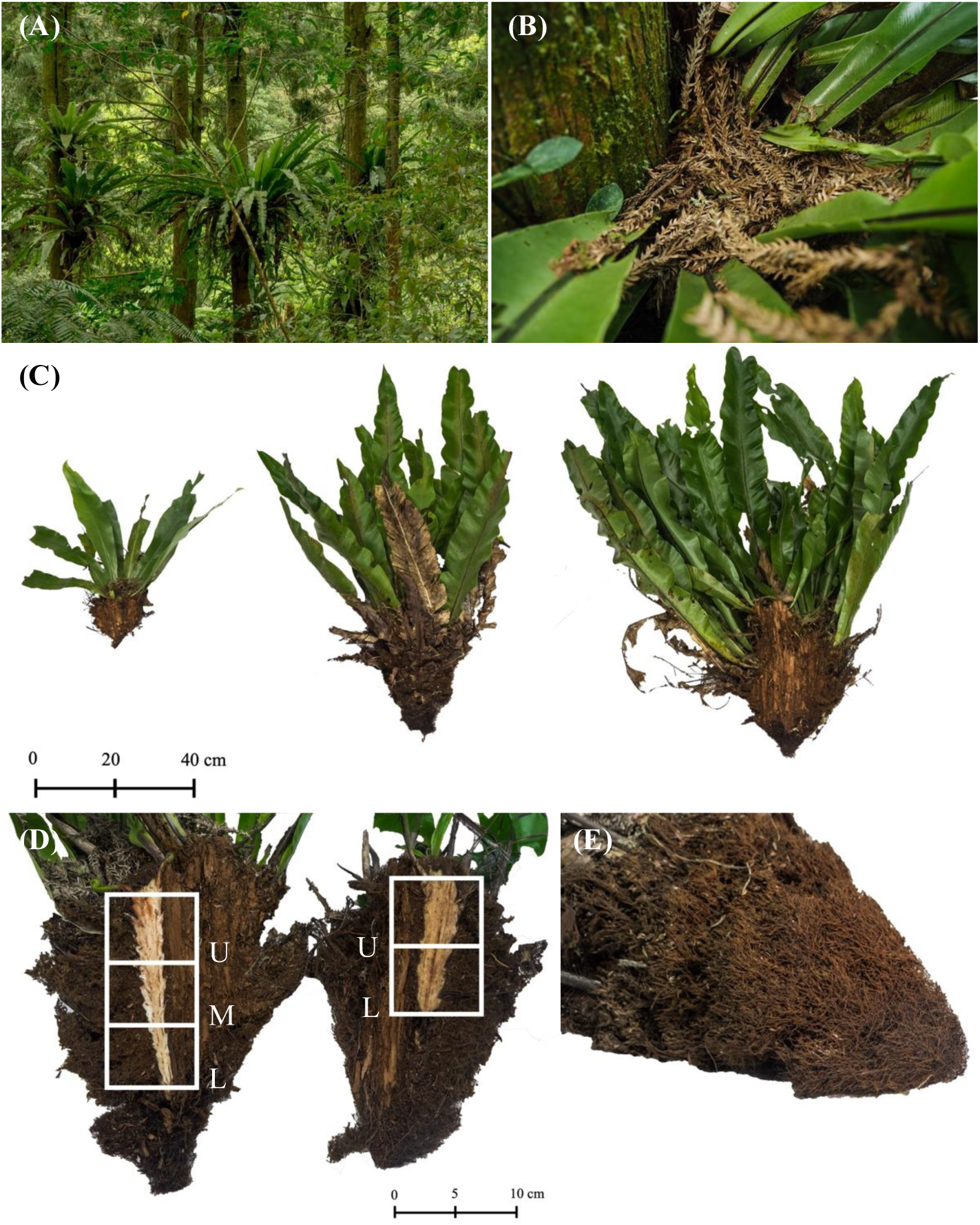
*Asplenium nidus* (bird’s nest fern; BNF) as a biotic island model system. (A) Study site showing BNF individuals growing in a *Cryptomeria japonica* plantation, selected to minimize large-scale environmental heterogeneity. (B) The litterfall from *Cryptomeria japonica* collected in a BNF nest structure, forming a suspended soil habitat. (C) BNF individuals were categorized into three size categories (from left to right) based on their rosette size: small (rosette size < 100 cm), medium (rosette size between 100 cm to 150 cm), and large (rosette size > 150 cm). (D) To collect the humus for DNA sequencing, a BNF individual was divided into three longitudinal slices; each slice was partitioned into two layers (upper (U) and lower (L)) for medium-sized BNF (right) and three layers (upper (U), middle (M), and lower (L)) for large BNF (left). (E) Additional humus sampling from the outer portion of medium and large BNF individuals.

### Sample collection

We classified BNF individuals into three size categories based on their rosette sizes, defined as the maximum diameter of the rosette with naturally hanging fronds: small (rosette size *<* 100 cm), medium (100 cm *≤* rosette size *<* 150 cm), and large (rosette size *≥* 150 cm; Fig. 1C). In total, 24 BNF individuals across the three size categories were sampled: 8 small, 7 medium, and 9 large. We ensured that no sampled BNF had another BNF growing above it, thereby avoiding interception of litterfall and stemflow by upper-positioned individuals. Before sampling, the spatial position of each BNF was recorded by measuring the horizontal distances among host trees and the attachment height of each fern, allowing us to estimate distances between BNFs using both two-dimensional and three-dimensional spatial coordinates. The maximum two-dimensional and three-dimensional distances were both around 40 meters. Each BNF was carefully removed from its host tree using a betel nut knife, with a mat placed on the ground to collect any dislodged litter. For morphological characteristics, three leaves were randomly selected from each individual to measure frond length. After measuring the length, width, and height of the nest structure, we separated the BNF individual into fronds, accumulated litter, and the nest structure and measured their wet mass separately. After humus collection (see below), the nest structure was air-dried, and its dry mass was recorded.

The humus within the nest structure was collected through a stratified sampling design to capture the heterogeneity within individual BNF and to avoid under-representing variation in larger BNFs. Each BNF was evenly divided into three longitudinal slices. For small BNFs, humus containing decomposed litter fragments and roots was randomly subsampled from each slice and pooled into a 15 mL Falcon tube. For medium and large BNFs, each slice was further partitioned into two (upper and lower) and three (upper, middle, and lower) equally sized layers along the longitudinal axis, respectively (Fig. 1D). Subsamples were taken from each layer within each slice and pooled by layer into a single 15-mL Falcon tube. In addition, humus from the outer adventitious roots covering the nest was collected (Fig. 1E), forming in total three and four different sampling locations for medium and large BNFs, respectively (65 samples in total).

To extract the humus, 5 mL of sterilized double-distilled water (*ddH*_2_*O*) was added to each tube, followed by ultrasonic bathing for 90 s. Litter fragments and roots were then transferred into a new 15-mL Falcon tube for a second washing step, after which these solid materials were discarded. The liquid fractions containing humus from both tubes were pooled and centrifuged at 14,000 rpm for 5 min. Finally, the supernatant was discarded and the humus pellet was stored at -20 ℃until further processing for DNA sequencing.

### Humus properties for environmental heterogeneity analysis

To characterize within-patch environmental heterogeneity, we measured humus chemical properties (C:N and pH) for each layer of a BNF individual. The remaining humus from each BNF layer was sieved through a 2-mm mesh and mixed to homogenize the sample. For C:N ratio, two 10 mg subsamples of sieved humus were taken from each layer as replicates and analyzed using the Elemental Analyzer (UNICUBE EA0002, Germany). For pH, 1 g of sieved humus was mixed with 10 mL of deionized water; humus–water suspensions were allowed to settle for at least one hour before we measured their pH using a glass electrode (CLEAN PH500, CLEAN Instruments Co. Ltd., Taiwan).

### DNA extraction and sequencing

Microbial DNA was extracted from 250 mg of humus samples using the DNeasy PowerSoil Pro Kit (Cat no. 47014; Qiagen, Germany) according to the manufacturer’s protocol. Library preparation and sequencing of both fungal and bacterial microbiomes were conducted by Tri-I Biotech Inc. (Taiwan). Amplicon libraries were prepared using two-step polymerase chain reaction (PCR), where the target DNA region was amplified in the first PCR (30 cycles) and the 8-mer barcodes were added for each sample in the second PCR (5 cycles). For fungal communities, the ITS1 region of fungal ribosomal DNA was amplified using primers ITS1 (5’–CTTGGTCATTTAGAGGAAGTAA– 3’) and ITS2 (5’–GCTGCGTTCTTCATCGATGC–3’). For bacterial communities, the V6–V8 region of the 16S rRNA gene was amplified using primers 968F (5’–AACGCGAAGAACCTTAC–3’) (Chen *et al*., 2011) and 1391R (5’–ACGGGCGGTGWGTRC–3’) (Jorgensen *et al*., 2012). PCR reactions (25 *µ*l) contained 2 ng template DNA, 12.5 *µ*l of KAPA HiFi polymerase (Roche), 0.75 *µ*l of both forward and reverse primer (each 10 *µ*M), and MQ water to a final volume of 25 *µ*l. Thermal cycling conditions were 94*^◦^*C for 5 min; 30 cycles of 94*^◦^*C for 30 s, 52*^◦^*C for 20 s, and 72*^◦^*C for 45 s; followed by a final extension at 72*^◦^*C for 10 min. Amplicons were verified on 2% agarose gels and subjected to a second PCR (5 cycles, identical cycling conditions) to incorporate single indices and Illumina adapters using 10 ng purified PCR product as template. Indexed products were purified, quantified fluorometrically (Qubit), pooled in equimolar concentrations, and libraries were prepared using the Celero^™^ DNA-Seq Library Preparation Kit (TECAN). Library concentrations were determined by qPCR, and sequencing was performed on an Illumina MiSeq platform (2 *×* 300 bp pairedend).

### Bioinformatic analysis

For fungal communities, the raw Illumina MiSeq ITS rDNA sequences were processed using the DADA2 ITS Pipeline Workflow (version 1.8; Callahan *et al*., 2016) in R (version 4.3.2 (R Core Team, 2021)), with R packages “dada2”, “ShortRead”, and “Biostrings”. Briefly, after primer removal, the sequences were filtered and trimmed using the filterAndTrim function with the following parameter setting: maxN = 0, maxEE = c(2, 2), truncQ = 2. Subsequently, paired-end reads were merged and chimeras were removed. Finally, the UNITE database was used for taxonomy assignment of each amplicon sequence variant (ASV) (Abarenkov *et al*., 2024).

For bacterial communities, the raw Illumina Miseq 16s rDNA sequences were processed using the Quantitative Insights Into Microbial Ecology 2 (QIIME 2) pipeline (version 2023.05). Sequences were first denoised using the DADA2 plugin (Callahan *et al*., 2016), which included quality filtering, read truncation (forward and reverse reads were truncated to 247 bp and 255 bp, respectively), and chimera removal. Finally, the classifier-consensus-vsearch plugin (Bokulich *et al*., 2018) was used for taxonomy assignment of ASVs against the SILVA NR128 99% 16S rRNA database (Quast *et al*., 2012, Yilmaz *et al*., 2014).

### Statistical analyses

All downstream analyses of microbial data were performed separately for fungal and bacterial composition data, using R packages “phyloseq” (McMurdie & Holmes, 2013), “vegan” (Oksanen *et al*., 2012), and “mobr” (McGlinn *et al*., 2019). The ASVs with fewer than five total reads across the entire dataset were removed from both the fungal and bacterial composition data. To examine the ISAR pattern, fungal and bacterial ASV counts from each sample (representing layer-level diversity) were first summed within each BNF to represent the total diversity on a biotic island. For all of the analyses, ASV count matrices for each BNF individual were rarefied to even sequencing depth (determined by the BNF with the lowest number of reads) separately for fungal and bacterial datasets, except for the multi-scale rarefaction analysis, where we followed the framework of Chase *et al*. (2019) (see below). Because richness was calculated from the rarefied, pooled community within each BNF individual, this measure represents rarefied gamma diversity in our study (i.e., *^γ^S_n_*; see Multi-scale rarefaction below). We considered the ISAR power model *S* = *cA^z^*, where *S* represents the ASV richness of the rarefied microbial community and *A* represents the dry mass of the nest structure (i.e., a proxy for size) of the BNF individual (Arrhenius, 1921). On a double-logarithmic plot, simple linear regression was applied to assess the relationship between log-transformed ASV richness on log-transformed dry mass of BNFs. The same analyses were repeated at different taxonomy levels to evaluate the consistency of the pattern.

To investigate the underlying mechanism of the observed ISAR, we calculated diversity metrics across multiple scales using ASV rarefaction curves constructed at different levels. Because the small BNFs did not contain clearly defined layers within the nest structure for stratified sampling, this analysis was restricted to medium and large BNFs. Specifically, we estimated diversity at the BNF individual level (*γ* diversity), layer level (*α* diversity), and the differences between the two levels (*β* diversity) following the framework of Chase *et al*. (2019). The BNF-level rarefaction curve (the *γ* rarefaction curve) was identical to that used in the aforementioned ISAR analysis, i.e., combining samples from all layers for each BNF prior to rarefaction. From this curve, we calculated *^γ^S_n_*, the expected number of ASVs observed when randomly sampling *n* individuals, where *n* was determined by the BNF individual with the lowest sequencing depth (Gotelli & Colwell, 2001). We also calculated *^γ^S_PIE_*, the effective number of species based on the probability of interspecific encounter (PIE, a measurement emphasizing common species), which was extracted as the slope of the base of the *γ* rarefaction curve. Finally, considering rare species, we estimated the total ASV richness (*S_total_*) from the raw (non-rarefied) abundance data using the Chao1 estimator (Chao, 1984, 1987). Alternatively, for each layer sample within a BNF individual, we constructed its layer-level rarefaction curve (the *α* rarefaction curve). We calculated the layer-level metric *^α^S_n_*, which represents the average expected number of ASVs observed when randomly sampling *n* individuals from the *α* rarefaction curve, where *n* was determined by the layer sample with the lowest sequencing depth (Gotelli & Colwell, 2001). We also quantified the corresponding *^α^S_PIE_*, which represents the average effective number of species across all layers within a BNF. For *β* diversity, which reflects compositional heterogeneity among layers within a BNF, diversity indices were calculated as ratios of gamma to alpha metrics: *^β^S_n_* = *^γ^S_n_*/*^α^S_n_*, *^β^S_PIE_* = *^γ^S_PIE_*/*^α^S_PIE_*. Here, the numerator represents the expected diversity from the *γ* rarefaction curve, whereas the denominator represents the average expected diversity from the *α* rarefaction curves. Simple linear regression was applied to investigate the relationship between these diversity indices and the dry mass of BNFs.

We interpreted the above linear regression following Chase *et al*. (2019). Under a pure passive sampling hypothesis, larger BNFs are expected to contain more ASVs solely because they harbor more individuals. Accordingly, after standardizing for the number of individuals sampled, *γ* diversity metrics (e.g., *^γ^S_n_*and *^γ^S_PIE_*) should not vary systematically with BNF dry mass. This implies that *γ* rarefaction curves should overlap across BNFs of different sizes. Alternatively, if *^γ^S_n_* or *^γ^S_PIE_* increased significantly with BNF dry mass, this indicates that the ISAR pattern cannot be explained by *passive sampling* alone. Additional positive relationships for the *α*-level indices suggest *disproportionate effects*, because higher rarefied diversity within layers on larger BNFs suggests that island size influences local ecological processes rather than simply increasing the number of individuals sampled. Finally, positive relationships at the *β*-level indices support a role for *environmental heterogeneity* in shaping the ISAR, as greater differences between *γ* and *α* diversity with increasing BNF size reflect greater species-assemblage differences among layers within larger BNFs (Chase *et al*., 2019).

To examine how environmental variability within BNFs relates to patch size and microbial diversity, we quantified variation in humus pH and C:N ratio, and related these measures to microbial dissimilarity among layers within each BNF. First, the environmental variability within each BNF was characterized by the ranges and variances of pH and C:N ratio across layers. These environmental variability metrics were then regressed against BNF dry mass to examine whether environmental heterogeneity correlates with BNF size. Next, microbial compositional dissimilarity within each BNF was quantified as the mean of pairwise Bray–Curtis distances among layers. Linear regression was then applied to assess the relationship between environmental variability and mean within-BNF microbial compositional dissimilarity, allowing us to assess whether environmental heterogeneity was associated with higher microbial diversity.

To test for the distance-decay pattern within the BNF insular system, we examined the relationship between the microbial compositional dissimilarity and the spatial distance among BNFs. Microbial compositional dissimilarities among BNFs were quantified as pairwise Bray– Curtis distances, using the rarefied BNF-level ASV community matrices as in the previous ISAR analyses. Euclidean spatial distances were computed between all pairs of BNFs to construct spatial distance matrices. Mantel tests were conducted to evaluate the correlations between microbial compositional dissimilarity and spatial distance for both fungal and bacterial communities. To account for potential spatial autocorrelation in environmental factors, partial Mantel tests were also performed with pH and C:N ratio included as conditioning variables, allowing us to assess whether environmental gradients influenced the observed distance-decay relationships.

## Results

We found a significant positive relationship between ASV richness (i.e., rarefied gamma diversity, *^γ^S_n_*) and dry mass of BNF for both fungal (Slope (*z*) = 0.30, *P* < 0.001, *R*^2^ = 0.75) and bacterial (Slope (*z*) = 0.35, *P* < 0.001, *R*^2^ = 0.59) communities (Fig. 2, Table S1). This positive species–area relationship was consistent across all taxonomic levels examined (Table S1), indicating that larger BNFs supported higher richness in both fungal and bacterial communities.

**Figure 2.**
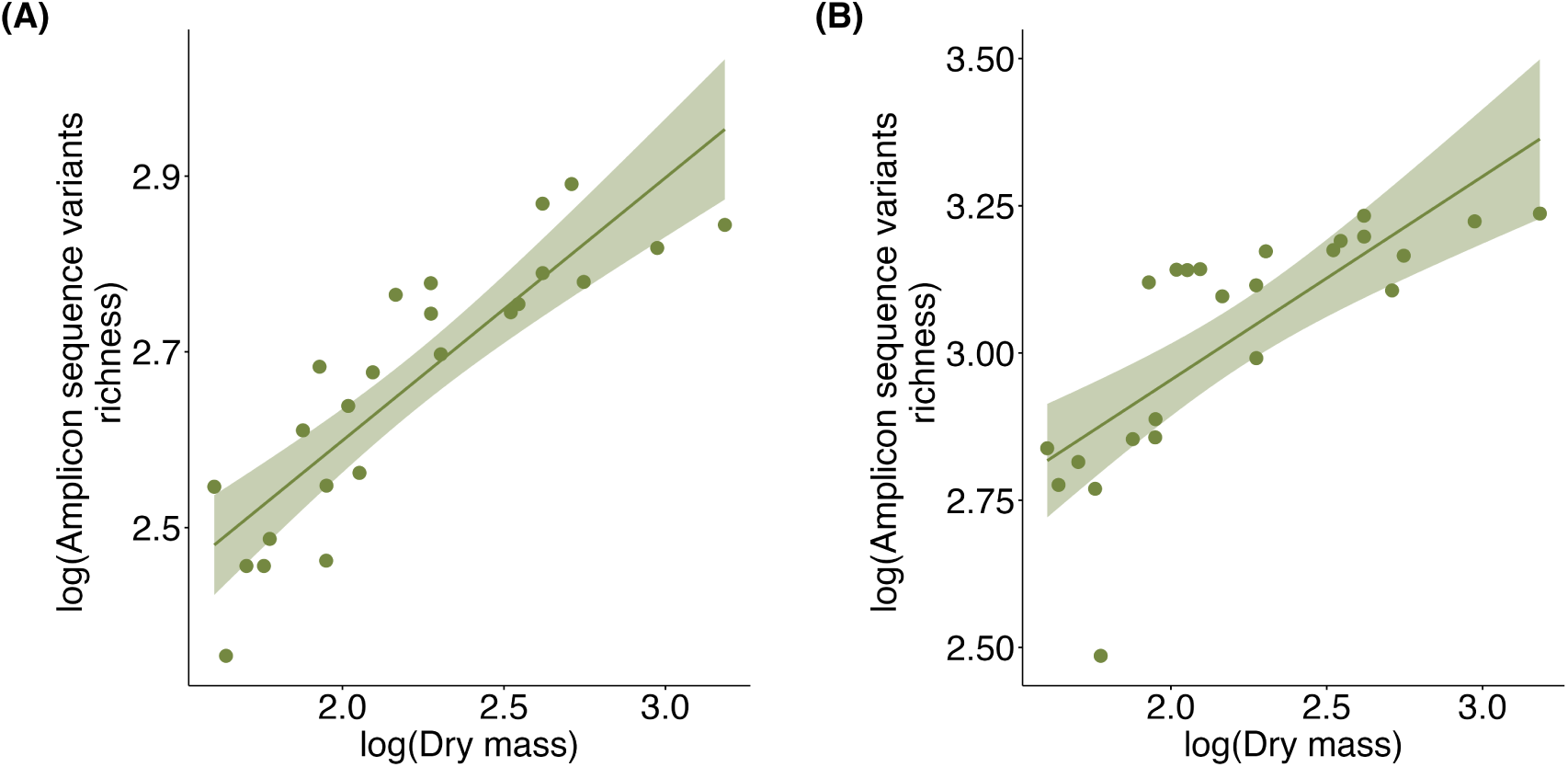
Significant positive relationship between amplicon sequence variant (ASV) richness and dry mass of *Asplenium nidus* (bird’s nest fern; BNF) individuals for (A) fungal communities (Slope(*z*) = 0.30, *P* < 0.001, *R*^2^ = 0.75) and (B) bacterial communities (Slope(*z*) = 0.35, *P* < 0.001, *R*^2^ = 0.59). Note the log-log scale on the axes.

Given the positive relationship between ASV richness of microbes and BNF dry mass, we next investigated the potential mechanisms underlying this pattern (Fig. 3, Table S2). The positive relationship between *S_total_*and the dry mass of BNF indicates that larger BNFs support higher observed richness, a common species–area pattern. However, because *S_total_*is not standardized for sampling effort, this pattern alone does not distinguish among potential mechanisms underlying the ISAR. Examining the standardized diversity metrics for the fungal community, both *^γ^S_n_* and *^γ^S_PIE_* increased significantly with the dry mass of BNF, rejecting the hypothesis that only *passive sampling* was operating in this system (Fig. 3A). Additionally, both alpha-level (*^α^S_n_*and *^α^S_PIE_*) and beta-level (*^β^S_n_*and *^β^S_PIE_*) indices showed significant positive relationships with BNF dry mass (Fig. 3B & C, Table S2). Together, these results suggest that both *disproportionate effects* and *environmental heterogeneity* contributed to shaping ISAR.

**Figure 3.**
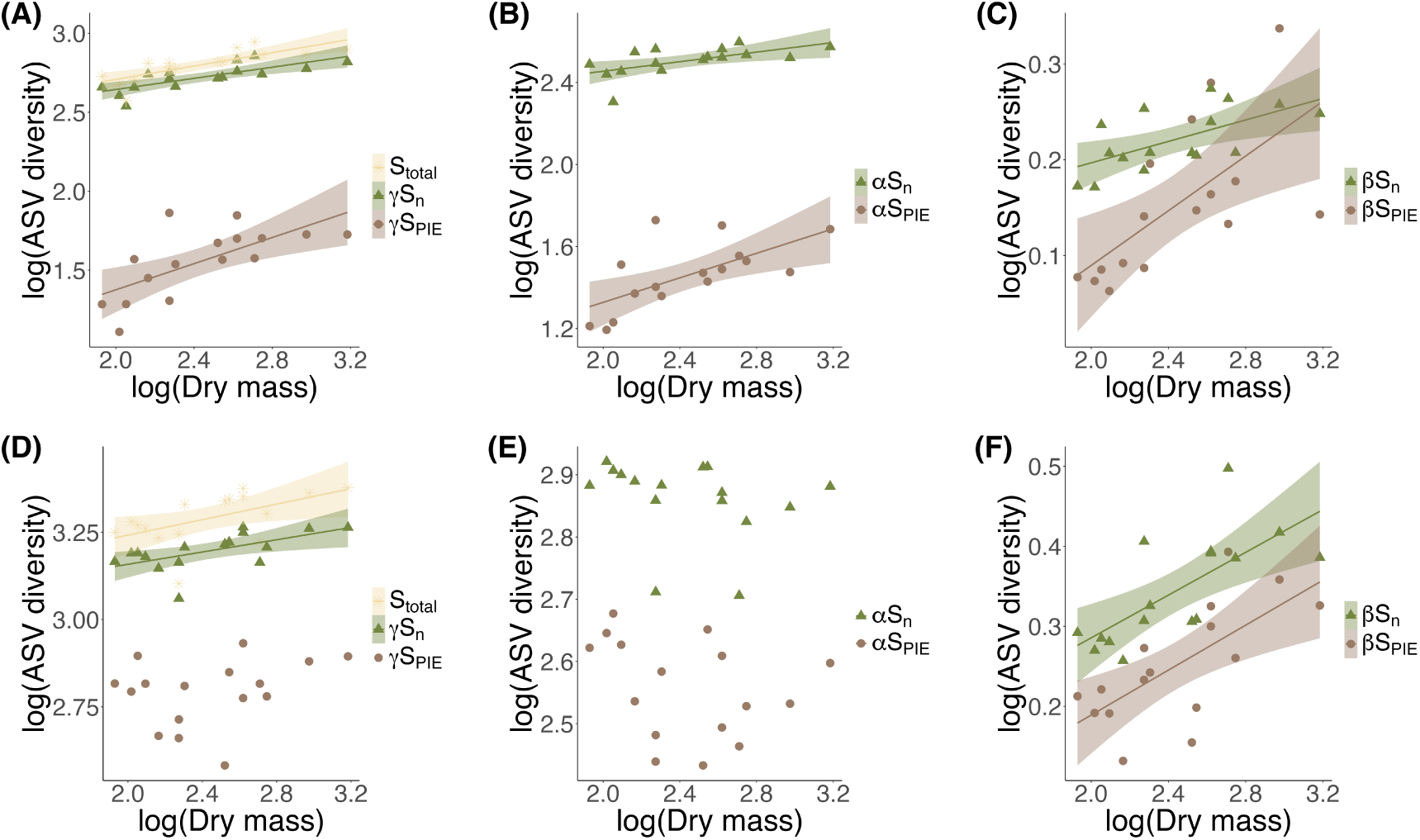
Relationships between ASV diversity indices at different scales and the dry mass of bird’s nest ferns (BNF) individuals for (A–C) Fungal and (D–F) Bacterial communities. Axes are shown on a log–log scale. (A & D) Diversity indices at the BNF level (*γ* rarefaction curves). *S_total_*represents the estimated ASV richness on a single BNF using the Chao1 estimator, which considers both rare and common ASVs; *^γ^S_n_* represents the expected number of species observed when randomly sampling *n* individuals while *^γ^S_PIE_* represents the effective number of species given the probability of interspecific encounter (PIE), a measurement of evenness that emphasizes common species. (B & E) Diversity indices at layer-level (*α* rarefaction curves). *^α^S_n_* represents the expected number of species observed when randomly sampled *n* individuals from a *α* rarefaction curve of a layer, whereas *^α^S_PIE_* represents the effective number of species across layers within a single BNF; both indices were averaged across all layers within each BNF individual. (C & F) Comparing diversity at the BNF individual level and the layer level. *^β^S_n_*considers both rare and common ASVs whiel *^β^S_PIE_* emphasizes more on common ASVs. Solid lines represent significant relationships, with statistic values shown in Table S2.

For the bacterial community, *^γ^Sn* increased with BNF dry mass while *S*PIE showed no corresponding trend, which may reflect an accumulation of additional, low-abundance ASVs in larger BNFs. This pattern indicates that mechanisms beyond *passive sampling* may have played a role in shaping ISAR (Fig. 3D). The relationship between *^α^S_n_* and BNF dry mass was not significant but *^β^S_n_* increased significantly with BNF dry mass, suggesting that *environmental heterogeneity*, rather than *disproportionate effect*, contributed to the ISAR pattern (Fig. 3E & F). Notably, neither *^γ^S_PIE_* nor *^α^S_PIE_* showed a significant relationship with the dry mass of BNF, indicating that the dominance structure of the community remained relatively constant across the BNF size gradient.

However, despite the lack of increases at the *α*- or *γ*-levels (i.e., without additional ASVs in larger BNFs), *^β^S_PIE_* demonstrated a significant positive relationship with the dry mass of BNF (Fig. 3F). This indicates that larger BNFs exhibit greater between-layer differentiation in the composition of common ASVs, potentially reflecting greater environmental heterogeneity that allows different taxa to dominate in different microhabitats.

Given the role of *environmental heterogeneity* in forming ISAR for both fungal and bacterial communities, we examined variation in pH and C:N ratio as potential contributors to heterogeneity within BNF. Among the measured environmental variables, only pH showed significant relationships with both BNF dry mass and microbial community dissimilarity among layers, whereas no significant patterns were detected for the C:N ratio (Fig. S2 and S3). We found that both the range of pH (*r* = 0.63, *P* = 0.009, Fig. 4A) and the variance of pH (*r* = 0.6, *P* = 0.014, Fig. S1A) within a BNF increased significantly with BNF dry mass. Additionally, the mean pairwise dissimilarity of both fungal and bacterial communities among layers within each BNF increased significantly with both pH range (fungi: *r* = 0.67, *P* = 0.004, Fig. 4B; bacteria: *r* = 0.59, *P* = 0.016, Fig. 4C) and pH variance (fungi: *r* = 0.65, *P* = 0.006, Fig. S1B; bacteria: *r* = 0.54, *P* = 0.032, Fig. S1C). These results suggest that increasing pH heterogeneity within larger BNFs is associated with greater microbial compositional turnover across layers, potentially resulting in the observed ISAR for both fungal and bacterial communities.

**Figure 4.**
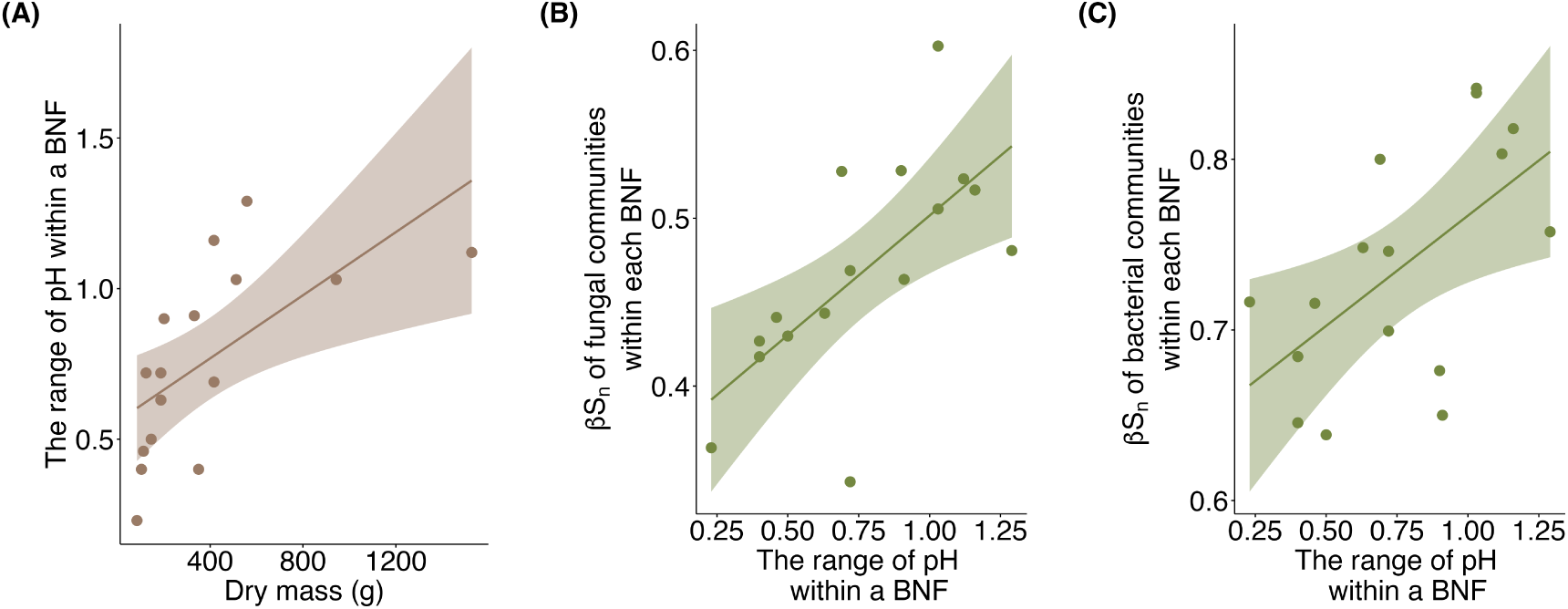
Environmental heterogeneity of pH as a potential mechanism for the observed species–area relationship. (A) Range of pH shows a significant positive relationship with BNF size (*r* = 0.63, *P* = 0.009). Mean pairwise dissimilarity of microbial communities across BNF layers increases with the pH range within a BNF individual; the positive relationship was significant for both (B) fungal (*r* = 0.67, *P* = 0.004) and (C) bacterial (*r* = 0.59, *P* = 0.016) communities.

Finally, regarding the spatial distance effect, ASV compositional dissimilarity increased with the spatial distance between pairs of BNFs for both fungal (Mantel *r* = 0.17, *P* = 0.025) and bacterial (Mantel *r* = 0.19, *P* = 0.017) communities (Fig. 5). When accounting for environmental variation using partial Mantel tests, controlling for pH did not alter the significant spatial relationship for either fungi (*r* = 0.17, *P* = 0.022) or bacteria (*r* = 0.15, *P* = 0.042). These results suggest that the distance-decay pattern of fungal communities is largely independent of measured environmental variables and likely reflects dispersal limitation. However, the spatial compositional dissimilarity of bacterial communities was slightly weaker after accounting for pH, suggesting a minor contribution of spatially structured environmental factors.

**Figure 5.**
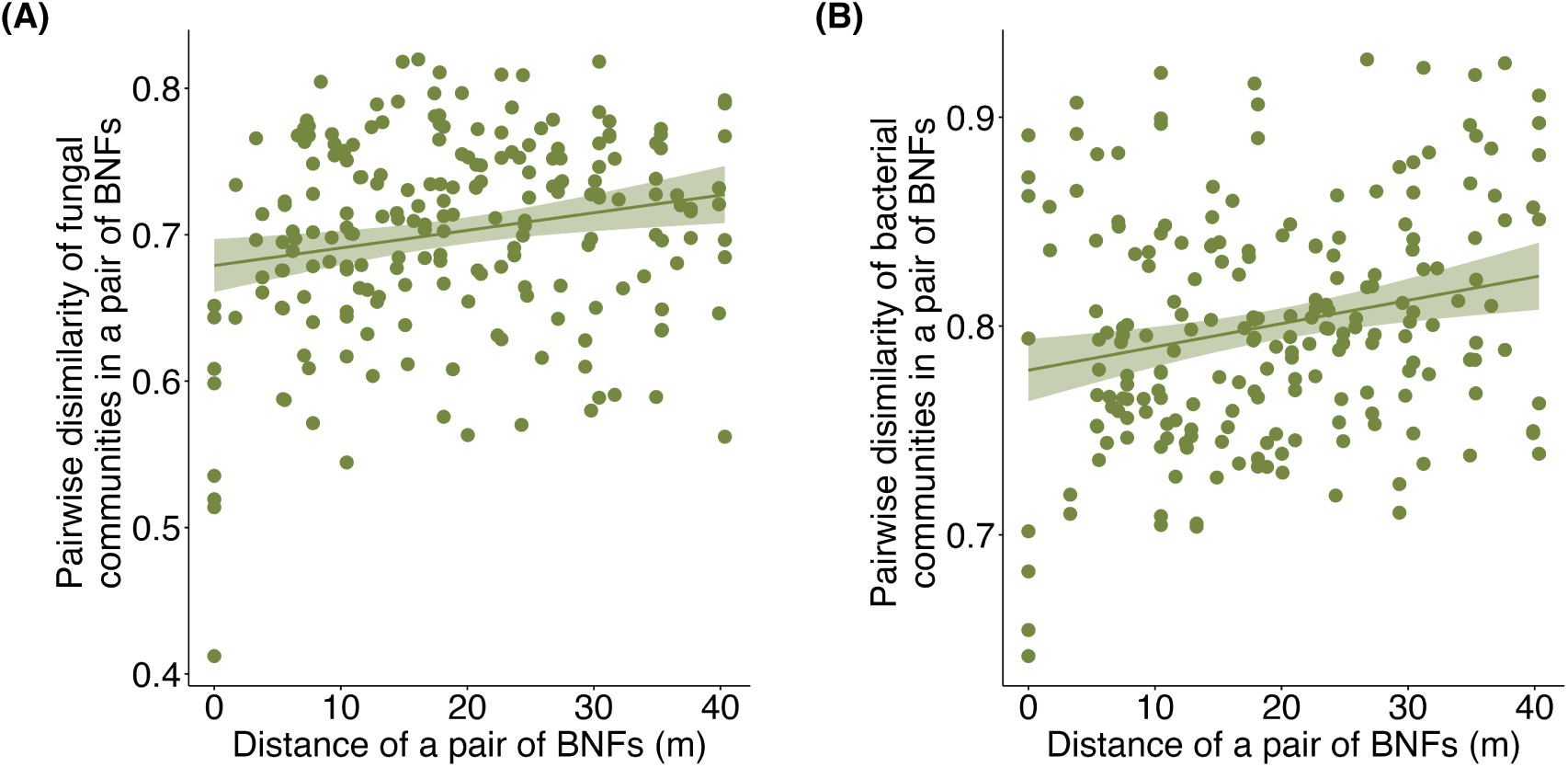
Relationship between ASV dissimilarity and the two-dimensional spatial distance among pairs of BNF individuals. The positive relationship was significant for both (A) fungal (Mantel *r* = 0.17, *P* = 0.025) and (B) In the bacterial (Mantel *r* = 0.19, *P* = 0.017) communities.

## Discussion

Consistent with our central hypothesis that organism-formed biotic islands conform to classical island biogeography predictions, both fungal and bacterial richness increased with BNF size (i.e., the species–area relationship). By evaluating the three mechanisms underlying ISAR, we found consistent support for *environmental heterogeneity* in both fungal and bacterial communities, whereas evidence for *disproportionate effects* was specific to fungal richness. Together, these results align with our hypothesis that organism-driven structural complexity within biotic insular systems would create more diverse microhabitats, thereby promoting microbial diversity.

Our results suggest that humus pH is a key environmental factor underlying the internal heterogeneity of BNFs. Larger BNFs exhibited greater pH variation among nest layers, particularly between the upper and lower layers of the nest structure, with differences of up to one pH unit in larger ferns. Importantly, this vertical pH gradient likely reflects the growth of the habitat-forming BNF: as individuals enlarge, newly intercepted litter accumulates in the upper layer, while older litter is retained below and progressively decomposes in lower layers. This process generates stratified microhabitats across layers and contributes to the environmental heterogeneity mechanism within this biotic insular system. This interpretation is consistent with the broader literature showing that soil and host-associated microbiomes are strongly structured by pH, with both bacterial and fungal community composition showing pronounced shifts along pH gradients (e.g., Hinsinger *et al*., 2003, Fierer & Jackson, 2006, Rousk *et al*., 2010, Bahram *et al*., 2018). Since both fungi and bacteria are sensitive to pH variation, the greater heterogeneity in larger BNFs likely expands the range of ecological niches and promotes microbial diversity. In some biotic insular systems, pH can be actively modified by the host, such as root-driven modification of the rhizosphere through proton extrusion and organic acid exudation (Hinsinger *et al*., 2003, Zhalnina *et al*., 2018), or host regulation along animal guts (Brinck *et al*., 2025). In contrast, pH heterogeneity in BNFs appears to emerge as the fern grows and retains litter across multiple stages of decomposition within the nest matrix. Together, these results indicate that environmental heterogeneity within BNFs is not merely a correlate of patch size, but a mechanistic pathway through which area influences microbial diversity in this biotic insular system.

The *disproportionate effect* detected for fungal communities likely reflects greater sensitivity of fungi to demographic constraints in smaller patches. This could be attributed to the fact that fungi are more dispersal-limited compared to bacteria (Li *et al*., 2020, Chen *et al*., 2020, Schmidt *et al*., 2014, Zhang *et al*., 2021), which may increase their vulnerability to stochastic extinction when population sizes are small. In contrast, bacteria typically exhibit smaller body and propagule sizes and better dispersal abilities (Farjalla *et al*., 2012, Powell *et al*., 2015), making them less susceptible to such demographic constraints and therefore less influenced by patch size. The contrasting responses of fungal and bacterial communities further suggest that trait differences between species may mediate how classical island biogeography mechanisms manifest in biotic islands.

Consistent with our hypothesis on isolation effects, microbial community similarity decreased with increasing projected distance between BNFs, demonstrating a clear distance-decay pattern. This pattern highlights the role of dispersal limitation as a regional process shaping community composition, consistent with predictions from the metacommunity framework, even over relatively short distances. Although both fungal and bacterial communities exhibited spatial structuring, this pattern contrasts with the inference from the ISAR mechanism analysis, in which evidence for *disproportionate effect*, and thus stronger indications for dispersal limitation, was observed primarily for fungi. The observed spatial pattern for bacteria may instead partially reflect spatially autocorrelated environmental factors, particularly pH, or other unmeasured variables that covary with distance and influence bacterial community assembly. Although distance-decay patterns are commonly reported at large geographic scales (Martiny *et al*., 2006, Hanson *et al*., 2012), recent studies indicate that dispersal limitation can also influence microbial community composition at much smaller spatial scales (Deakin *et al*., 2018), as observed in the BNF system, where microbial dispersal is restricted even among nearby nests.

A key challenge in studying organism-formed habitat patches is the confounding relationship between patch area and organism age, both of which can influence the diversity of resident communities (Jones *et al*., 1994, Whittaker *et al*., 2007). This issue is a common feature of biotic insular systems, in which larger patches often correspond to older individuals that have had more time to accumulate resources, creating a natural correlation between area and age. Unlike abiotic islands, such habitats are generated and maintained by living organisms and their developmental dynamics, introducing additional dynamics in how area, age, and isolation shape resident diversity. In this study, we used dry mass as a proxy for patch area; however, this metric is inherently correlated with age because older ferns tend to accumulate more litter and therefore attain greater mass. This natural phenomenon limits our ability to fully disentangle the independent contributions of area and age on microbial diversity. Unlike woody plants, BNFs lack distinct age markers such as tree rings, making age estimation difficult. Future studies could address this limitation through long-term monitoring of BNF individuals or greenhouse experiments with BNFs of known age, which may allow clearer separation of area- and age-related effects on community assembly processes.

We argue that BNFs provide a promising model for studying microbial biodiversity in biotic insular systems. In this study, we controlled key factors such as host tree species and litter composition, creating a relatively homogeneous environment that allowed us to isolate the effects of dispersal and community assembly processes. Beyond this controlled setting, however, BNFs in complex canopy environments offer opportunities to explore a broader range of ecological questions. For example, BNFs growing in mixed-species forests (Fig. 6A) may receive litter inputs with variable litter chemistry (Binkley & Giardina, 1998) and co-occur with other epiphytic plant species (Fig. 6B), both of which may introduce patch-level environmental variability via plant-mediated nutrient cycling (Berendse, 1994) or plant–microbe interactions (Bever *et al*., 1997). Such variation would make BNFs a useful system for examining species sorting in addition to dispersal effects. Another dimension of interest is the vertical spatial structure of BNFs, where individuals are distributed both vertically and horizontally in a simple forest (Fig. 6C). Such stratification may influence nutrient flow and dispersal pathways, generating interconnected local communities (Tatsumi *et al*., 2021) for studying how spatial configuration alone can drive metacommunity dynamics. By leveraging their natural variability and discrete patch structure, BNFs offer a tractable system for future studies to investigate how multi-scale processes interact to shape microbial communities in complex, organism-formed insular habitats.

**Figure 6.**
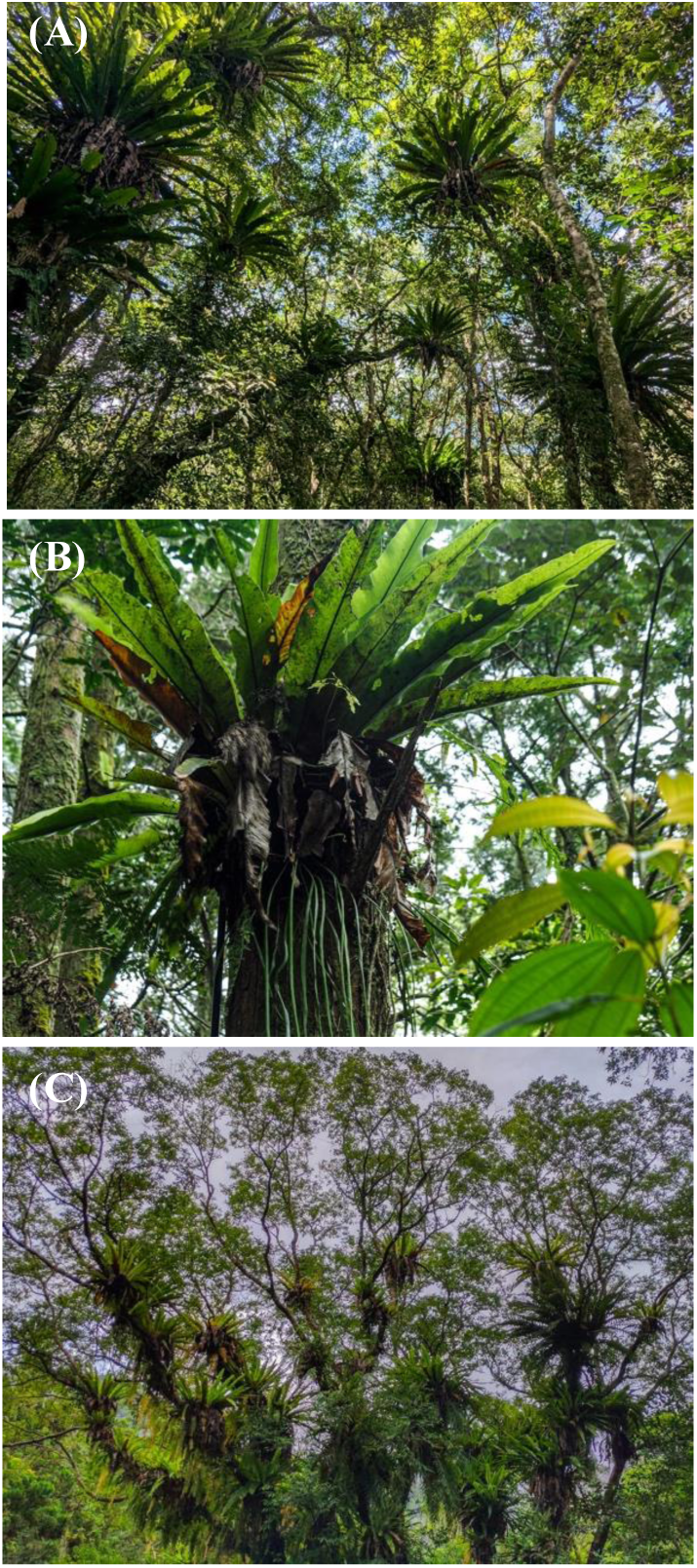
The spatial structure and environmental variability of bird’s nest ferns (BNFs) creating a naturally replicated metacommunity system, suitable for exploring multiscale community assembly processes. (A) BNFs growing in a mixed-species where variation in litter chemistry introduces patch-level environmental heterogeneity. (B) BNFs hosting other plant species, generating additional environmental variability through plant–microbe interactions and nutrient modification by co-occurring plants. (C) Vertical and horizontal distribution of BNFs in a pure forest. BNFs form interconnected habitat patches that enable the investigation of nutrient flows and species interactions across spatial layers.

In conclusion, our findings demonstrate that organism-formed patches can function as “islands”, in which both patch size and spatial isolation structure species diversity within the biotic insular systems. In the BNF system, patch growth is associated with litter accumulation and the development of pronounced decomposition gradients, thereby linking local habitat diversification within the patch-forming organism to microbial diversity patterns. Overall, our results demonstrate that BNFs, as naturally replicated biotic islands, can be used not only to study fine-scale assembly mechanisms but also to test and validate core predictions of classical island biogeography theory at spatial scales relevant to regional dispersal and patch connectivity. Importantly, the underlying mechanisms in this system are generated dynamically through organism-mediated habitat construction rather than imposed by static landscape geometry. We argue that by integrating organism-mediated habitat complexity with regional dispersal constraints, biotic insular systems such as BNFs provide a powerful system for extending island biogeography and metacommunity theory to self-modifying landscapes.

## Acknowledgements

We thank Yi Sun, I-Chih Shia, Hsun-Hung Chu, Ching-Ning Yeh for assisting with data collection. We thank Joe Wan, Niv DeMalach and members of the Ke Lab for providing feedback on the manuscript. This study is supported by the Yushan Fellow Program, Ministry of Education, Taiwan (MOE-110-YSFAG-0003-001-P1).

## Author contributions

PJK, YPT, and SW conceived the study; YPT and SW conducted field work; YPT analyzed the data and wrote the first draft of the manuscript; all authors contributed to the final version of the manuscript.

## Data accessibility statement

Should the manuscript be accepted, all primary data and analysis scripts will be archived in a public repository with the DOI included at the end of the article.

## Supplementary information

### Supplementary Tables and Figures

**Table S1.**
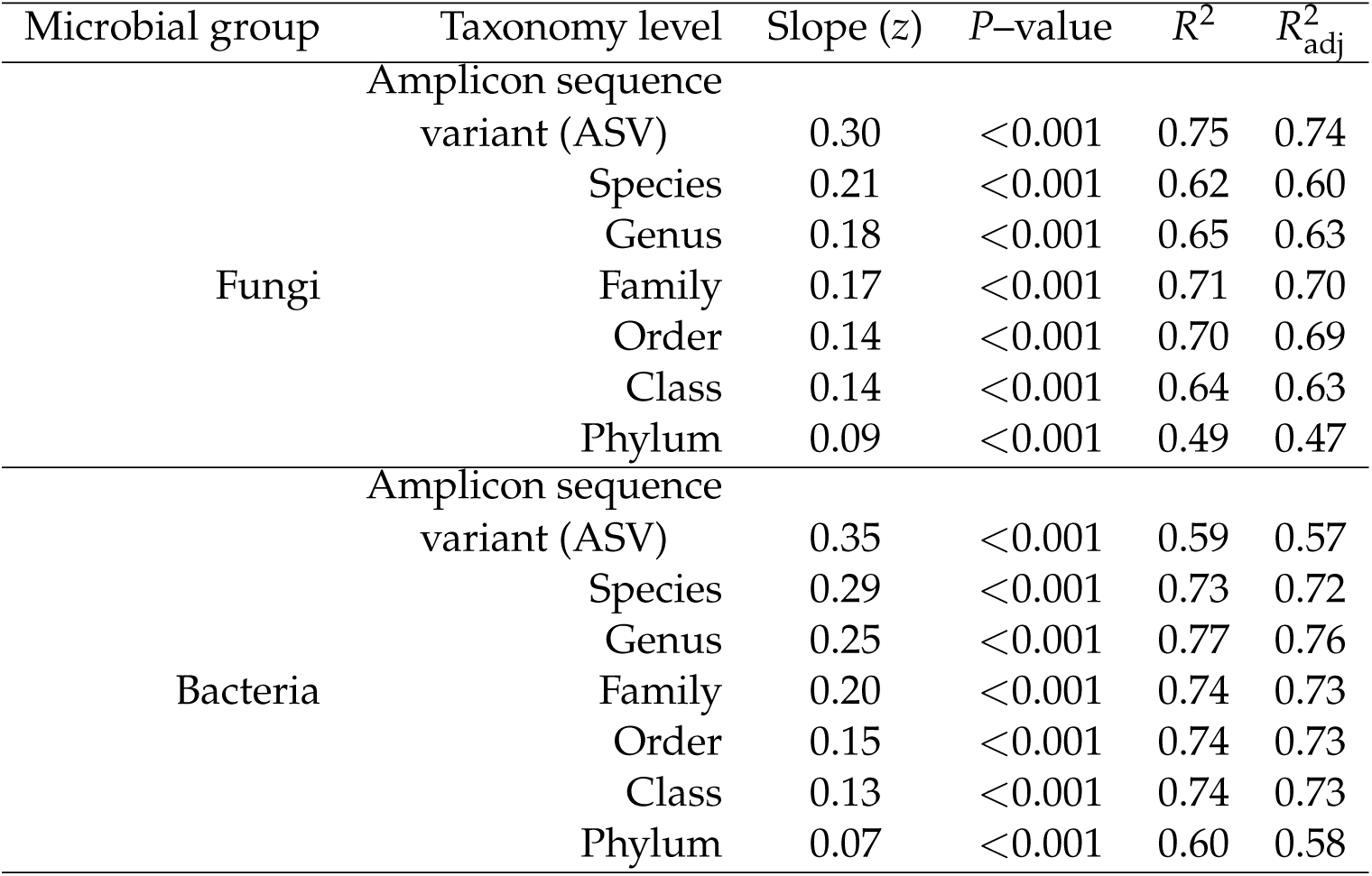
The results for simple linear regression with richness calculated based on different taxonomy levels for fungal and bacterial communities regressed on the dry mass of BNF. A significant positive relationship was shown across all the taxonomy levels for both fungal and bacterial communities.

**Table S2.**
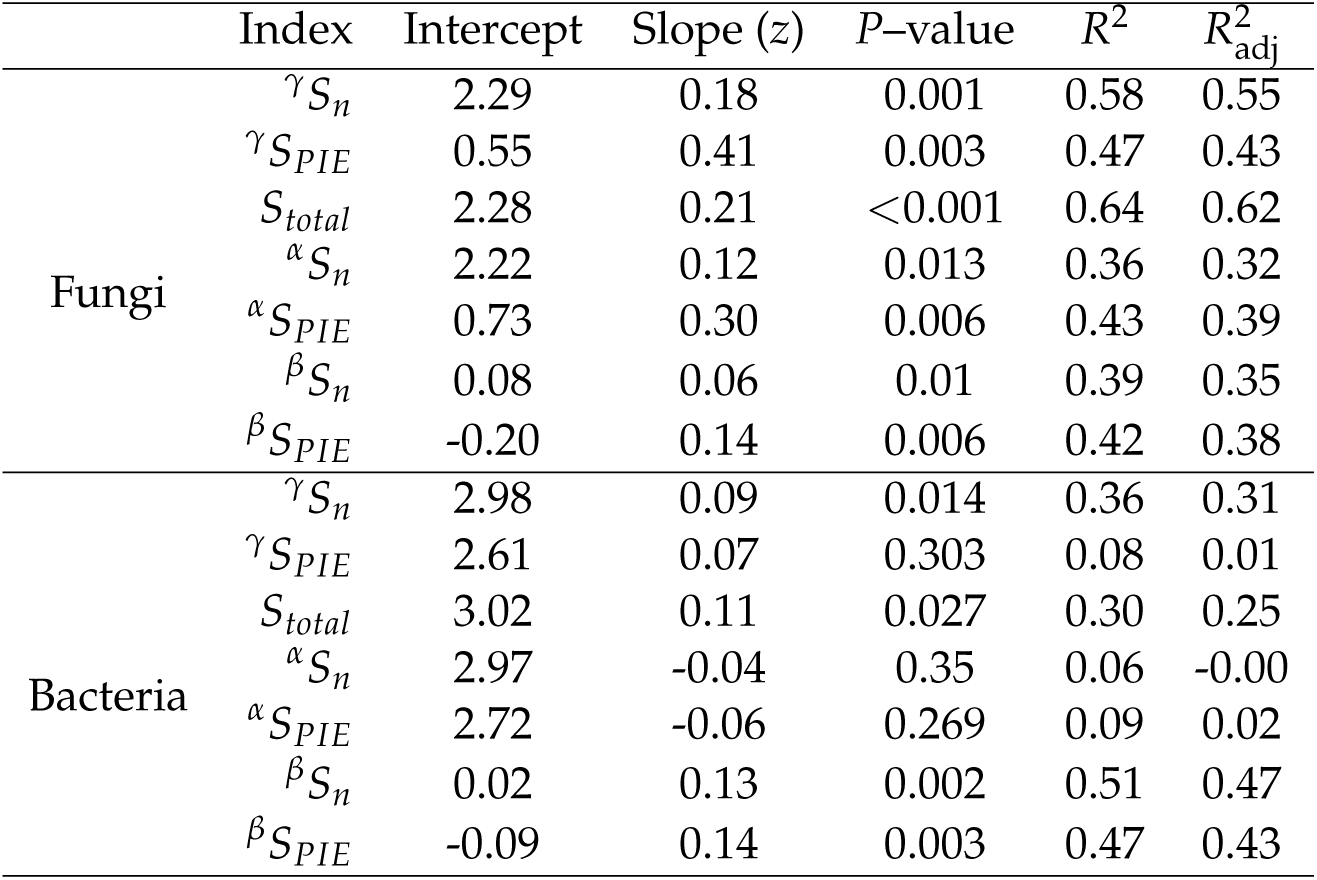
The results for simple linear regression with the diversity index at different scales by using the rarefaction curve for fungal and bacterial communities regressed on the dry mass of BNF in log scale.

**Figure S1.**
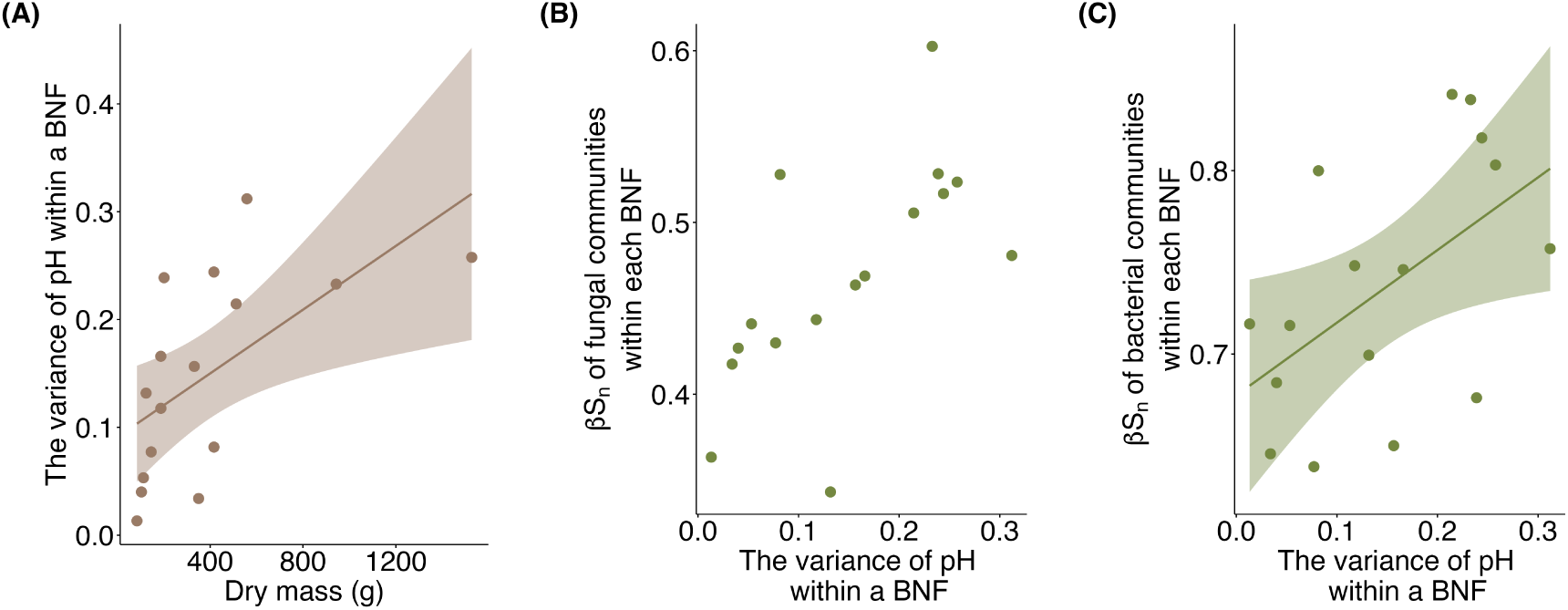
(A) Variance of pH shows a significant positive relationship with the size of bird’s nest ferns (BNFs) (*r* = 0.60, *P* = 0.014). (B) Mean pairwise dissimilarity of fungal ASVs across BNF layers increases significantly with pH variance (*r* = 0.65, *P* = 0.006). (C) Mean pairwise dissimilarity of bacterial ASVs across BNF layers increases significantly with pH variance (*r* = 0.54, *P* = 0.032).

**Figure S2.**
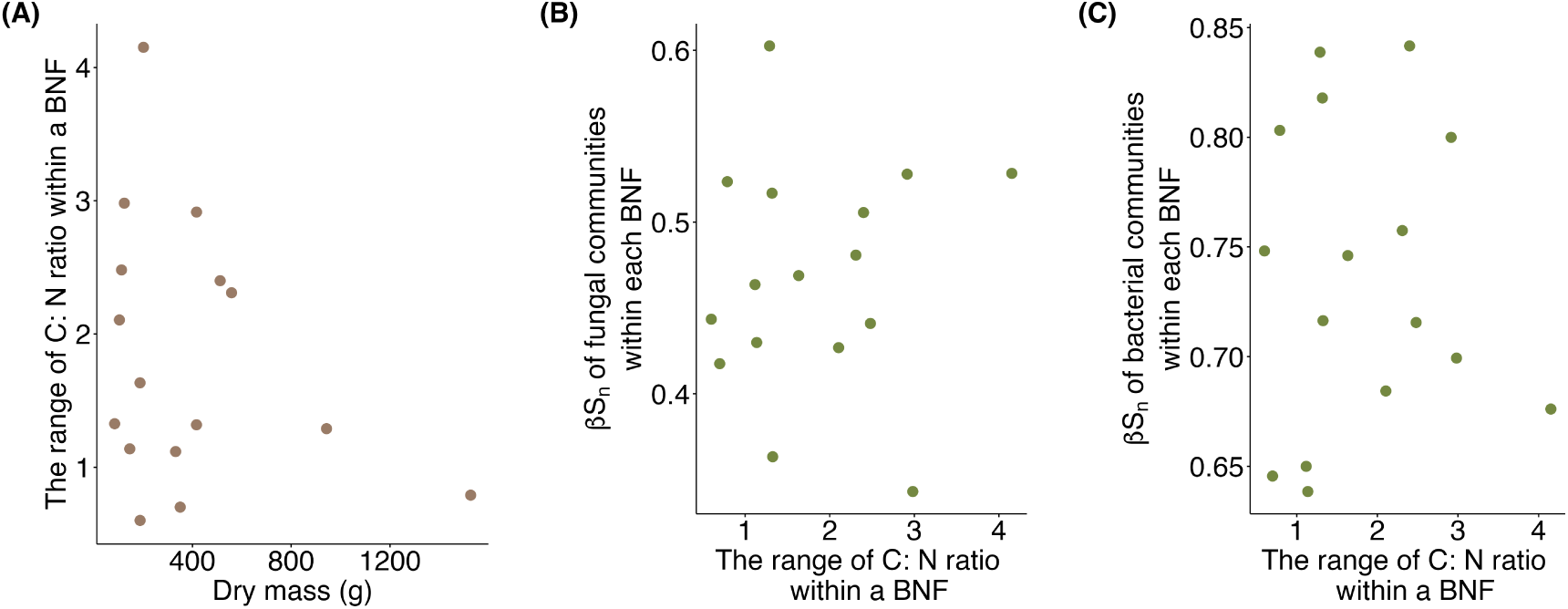
(A) Range of C:N shows no significant relationship with BNF size (*P* = 0.277). (B) Mean pairwise dissimilarity of fungal ASVs across BNF layers shows no significant relationship with C:N range (*P* = 0.783). (C) Mean pairwise dissimilarity of bacterial ASVs across BNF layers shows no significant relationship with C:N range (*P* = 0.964).

**Figure S3.**
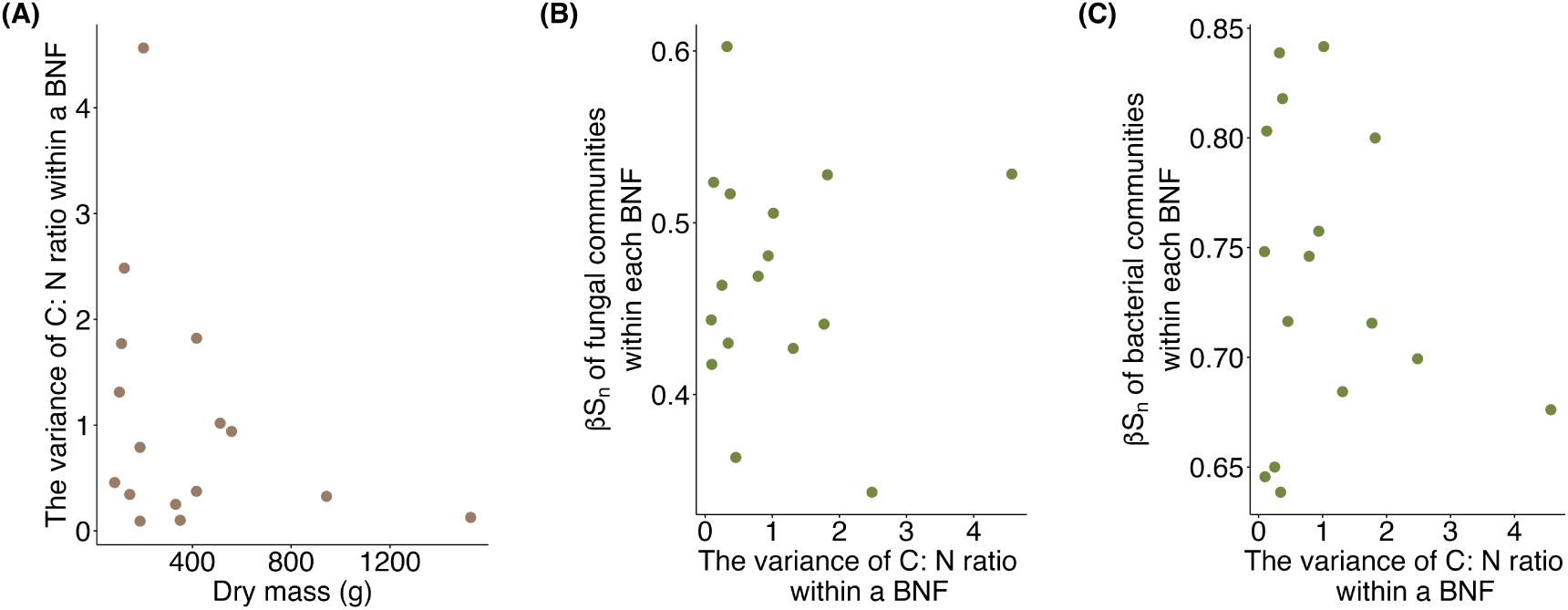
(A) Variance of C:N shows no significant relationship with BNF size (*P* = 0.235). (B) Mean pairwise dissimilarity of fungal ASVs across BNF layers shows no significant relationship with C:N variance (*P* = 0.929). (C) Mean pairwise dissimilarity of bacterial ASVs across BNF layers shows no significant relationship with C:N variance (*P* = 0.505).

